# Molecular Mechanisms for Frequency Specificity in a *Drosophila* Hearing Organ

**DOI:** 10.1101/2021.07.11.451973

**Authors:** Yufei Hu, Yinjun Jia, Tuantuan Deng, Ting Liu, Wei Zhang

## Abstract

Discrimination for sound frequency is essential for auditory communications in animals. Here, by combining *in vivo* calcium imaging and behavioral assay, we found that *Drosophila* larvae can sense a wide range of sound frequency and the behavioral specificity is mediated with the selectivity of the lch5 chordotonal organ neurons to sounds that forms a combinatorial coding of frequency. We also disclosed that Brivido1 (Brv1) and Piezo-like (Pzl), each expresses in a subset of lch5 neurons and mediate hearing sensation to certain frequency ranges. Intriguingly, mouse Piezo2 can rescue *pzl*-mutant’s phenotypes, suggesting a conserved role of the Piezo family proteins in high-frequency hearing.

## Introduction

Frequency discrimination of sound is a prerequisite to decode auditory signals (1, 2). In flies, frequencies of attractive conspecific courtship songs are distinct from those made by predators, and the auditory features can be unambiguously decoded by hearing organs (3). Like many other insect species, flies hear with non-tympanal ears (2, 4). Johnston’s Organ (JO) in the antenna, as a specialized Chordotonal organ (Cho), are their main hearing organs (5). Frequency specificity has been observed in the JO (6, 7), demonstrating the critical role of frequency coding of peripheral auditory neurons. However, how the tonotopical gradient is established is unknown.

It’s thought that flies’ hearing organs are optimally toned to sound frequency at hundreds Hz, as the courtship song and sounds from some prominent predators fall into this range (4, 8). However, *Drosophila* is also reported to be a prey of other animals in the wild, such as swallow or bats (9–11). Distinct from low frequency sounds (around 500 Hz) made by wasps (12), songs made by birds are often with higher frequencies (13). It’s still an open question whether flies are able to hear higher frequency sounds above 1 kHz.

The tonotopy of auditory sensory neurons maybe generated by both structural and molecular gradients(14, 15). Several transient receptor potential (TRP) channels have been well studied for their roles in hearing transduction in *Drosophila* (12, 16, 17), including Nanchung (Nan), Inactive (Iav) and No mechanoreceptor potential C (NompC)(18, 19). However, those proteins are expressed in all five neurons of lch5(20–22) and the mutants of these gene caused non-selective hearing defects (Figure. S3A). So far, no molecule has been reported with functional tonotopy in Cho neurons. In fly larvae, a group of neurons in the body wall, lateral chordotonal 5 (lch5) neurons are the major sensors for vibration and sound (12). We thus speculated that genes with differential expression patterns among lch5 neurons may contribute to the heterogeneity of their frequency response.

In the present study, by combining *in vivo* calcium imaging and behavioral assay, we demonstrate that *Drosophila* larvae can sense a wide range of sound frequencies and the lch5 neurons mediate tonotopy. We also evaluated the distinct roles of Brivido1 (Brv1) and Piezo-like (Pzl) in hearing sensation and found they function at different hearing frequencies. The high frequency hearing defect of *pzl* mutant could be rescued by expressing mouse Piezo2 in Cho neurons, suggesting a functional conservation between *Drosophila* Pzl and mammalian Piezo2.

## Results

### *Drosophila* larvae could sense high frequency sound

Sounds made by predators are often alerts to prey. Distinct from low frequency sounds (around 500 Hz) made by wasps(12), songs from bats or birds are often of high frequencies(13). To identify whether *Drosophila* larvae could sense high frequency sound, wild type larvae were stimulated with a bird song from the meadow bunting (*Emberiza cioides*) (Audio 1). After a 90-s stimulation, most of the wild type larvae ran away from the sound source, indicating that the sound caused aversive auditory response (Figures 1A and 1D). Spectral analysis of the bird song revealed the main frequencies were 4,003 Hz, 4,912 Hz and 7,296 Hz (Figures 1B and S1). To better define the stimulation, an artificial composite tone with an equal power combination of 4 kHz, 5 kHz and 7.3 kHz was used for the avoidance behavior assay. Not only the composite tone, but also the pure tones from each three frequencies all led wild type larvae run away (Figures 1C and 1D). These suggest that wild type larvae could sense high frequency sound.

**Figure 1.**
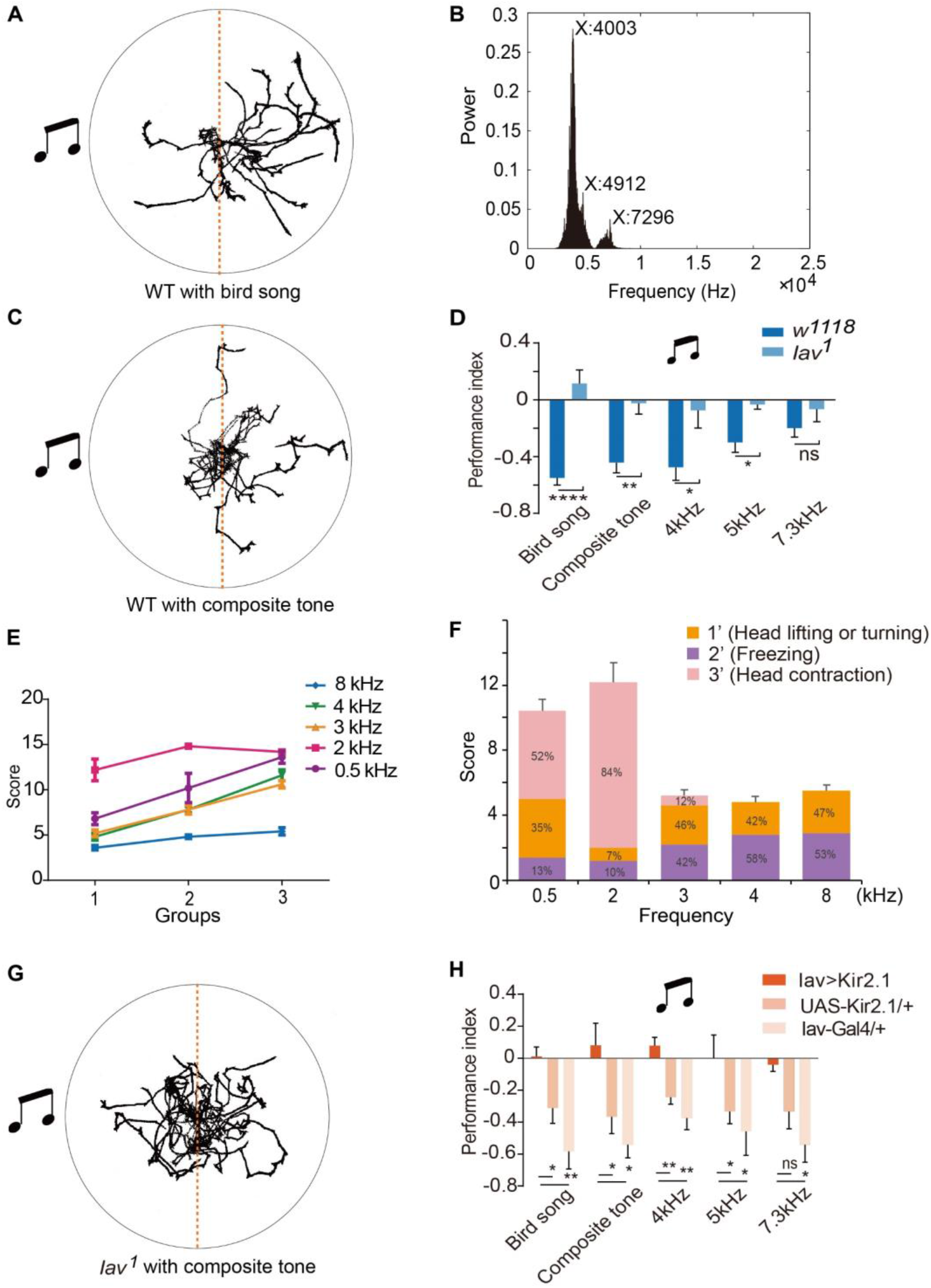
*Drosophila* larvae could sense high frequency sound. See also Figure S1, Movie S1 and Audio S1. (A) An example of avoidance behavior for wild-type larvae (*w*^*1118*^, n=20) with a bird song (90 s, 94 dB, SPL, sound pressure levels, Audio S1) stimulation. (B) Bird song frequency analyses. (C) An example of avoidance behavior for wild-type larvae (*w*^*1118*^, n=20) with a composite tone stimulation (90 s, 94 dB, SPL). (D) Performance index analyses for wild-type larvae (*w*^*1118*^) stimulated with sounds (bird song, composite tone, 4 kHz, 5 kHz and 7 kHz) for 90 s and an intensity of 94 dB (SPL, 20 larvae for each group and n=5-8, mean ± SEM, p*<0.05, p**<0.01, p****<0.0001 and ns, no significance, unpaired t test). (E) Single third instar larva was stimulated with different frequencies of sound with different intensities (intensities of 0.5, 2, 3, 4 and 8 kHz are divided into three groups with 5 dB increment: 80, 92, 91, 90, 84 dB, SPL for group1, 85, 97, 96, 95, 89 dB, SPL for group2, and 90.2, 102, 101, 100, 94 dB, SPL for group3; n=5). (F) Sound responses and the proportions of each score of wild-type larvae stimulated with different frequencies of sound with an intensity of 93 dB (SPL, n=5). (G) An example of avoidance behavior for *iav^1^* larvae (n=20) with a composite tone stimulation (90 s, 94 dB, SPL). (H) Performance index analyses for larvae with Cho neuron blocked (iav-Gal4/UAS-Kir2.1). And each stimulation lasted for 90 s with an intensity of 94 dB (SPL, 20 larvae for each group and n=5, mean ± SEM, ns, no significance, * for p<0.05, ** for p<0.01, *** for p<0.001 and **** for p<0.0001, one-way ANOWA with Dunnett’s multiple-comparisons test).

A freely moving larva usually explore an open field with a rhythmic locomotion pattern. Upon stimulation by the bird song, it displayed immediate head contraction, freezing, head lifting or turning (Movie S1). To quantify the responses, we applied the following scoring criterion: 1 point for head lifting or turning, 2 for freezing and 3 for head contraction. Each larva was stimulated with a 1-s sound with an interval of 10 s for 5 times at different frequencies (0.5, 2, 3, 4, and 8 kHz). The response score increased with the sound intensity (Figure 1E). The response patterns differed at different sound frequencies. Compared with higher frequencies, larvae performed more head contraction behavior when given lower frequency sound (Figure 1F).

Iav functions as a part of the mechano-electrical transduction (MET) machinery in Cho neurons, and flies with a loss function of *iav* were found insensitive to low frequency sound sensation(20). To explore whether Iav is also involved in high frequency sensation, *iav* mutant larvae were stimulated with the bird song, the composite tone, 4 kHz, 5 kHz or 7 kHz pure tone, respectively. No significant avoidance behavior was observed (Figures 1D and 1G). These results indicate that the *iav* mediated mechanotransduction is essential for the high frequency sound response.

### Five lch5 neurons show distinct frequency discrimination

Previous studies indicated that Cho neurons are responsible for vibration sensation in *Drosophila* larvae, a function dependent on Iav (12). Given that *iav* mutant larvae were defective in sensing high frequency sound (Figure 1D), we wondered whether Cho neurons are also required for high frequency sensation. Larvae with Iav^+^ neurons blocked showed no significant avoidance when stimulated with the bird song, composite tune or the three pure tunes (Figure 1H). This result suggests that Cho neurons are also involved in high frequency sensation.

To understand the neural mechanisms for high-frequency hearing in the fly larvae, we performed *in vivo* calcium imaging to characterize individual Cho neurons to various frequencies of sounds. By expressing Ca^2+^ indicator GCaMP6f (23) with Cha-Gal4 (24), the calcium responses of Cho neurons were monitored. The dendrites and cell bodies of the lch5 neurons responded to the stimulation within 1 second. Each lch5 neuron showed different response pattern to the frequencies tested (Figure 2A). The 1^st^ lch5 neuron was more sensitive to lower frequencies under 3 kHz, while the other four neurons were activated by sounds with frequencies higher than 0.5 kHz. The 2^nd^, 4^th^ and 5^th^ lch5 neurons responded broadly while the 3^rd^ lch5 neuron was only involved in 5 kHz sound sensation in our experimental setting (Figures 2B–2G). To rule out the possibility that high-frequency sound may cause low-frequency vibration of the substrate, we carried out non-contact vibration analysis and found that the glass slide frequency did not vibrate significantly upon stimulation of 5 kHz pure sound stimulation (Figure S2). This indicates that frequency discrimination is mediated by distinct pattern of lch5 neurons activated in *Drosophila* larvae.

**Figure 2.**
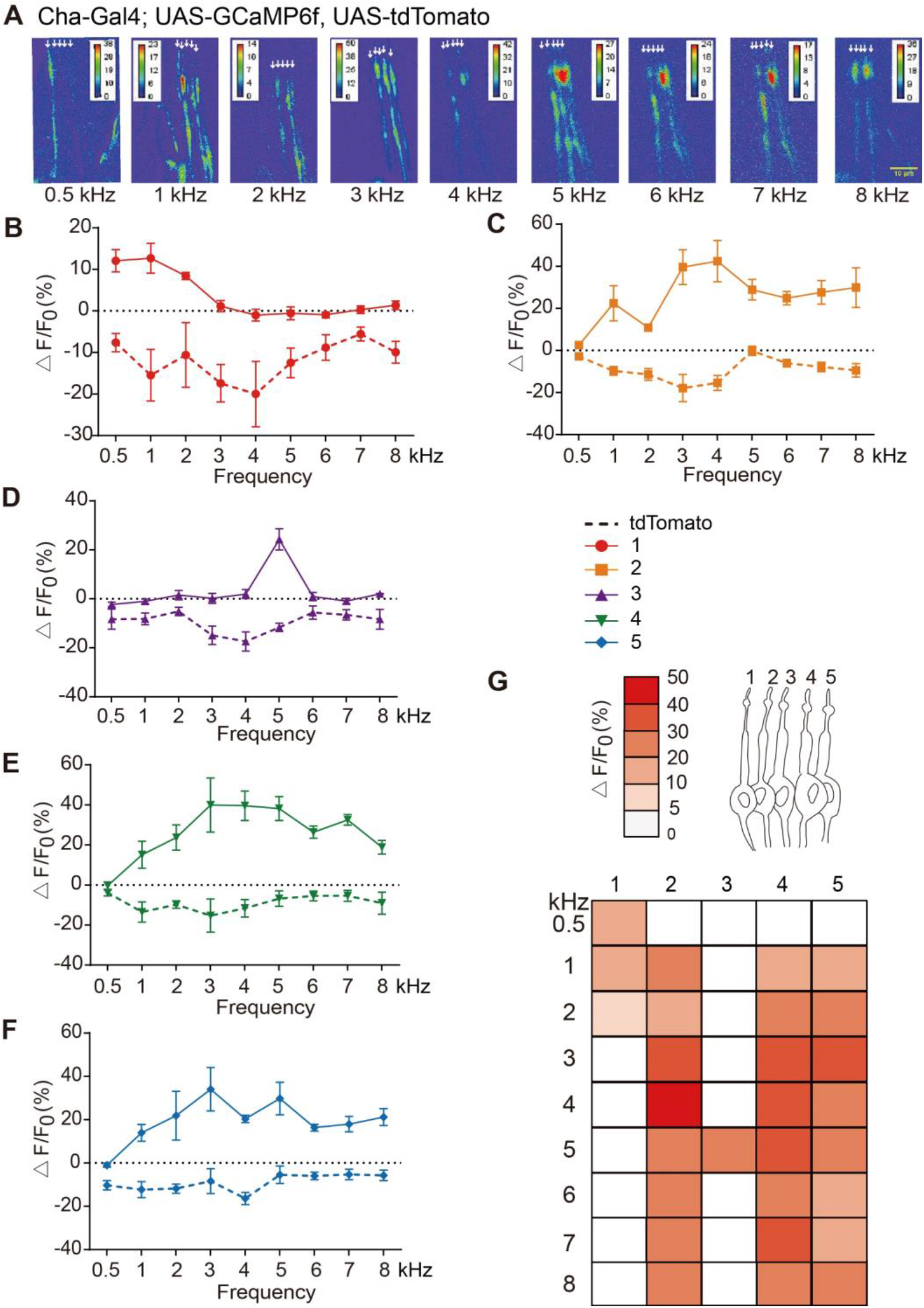
Five lch5 neurons show distinct frequency discrimination. See also Figure S2. (A) Stimulation by different frequencies of sound induced Ca^2+^ levels change in the dendritic tips of lch5 neurons. Arrows indicate the five neurons. (B-F) Statistical analysis of the Ca^2+^ responses of each lch5 neuron (1 to 5) to different sound stimulations in wild-type larvae. The region of interests (ROIs) were selected with a same area below cilium of each neuron (Cha-Gal4, UAS-GCaMP6f; UAS-tdTomato, mean ± SEM, n=5). (G) Diagrams for lch5 neurons (upper) and Ca^2+^ level changes in each lch5 neuron under different sound stimulation (lower).

### Brv1 and Pzl mediate hearing sensation to certain frequency ranges

To explore the molecular basis for the frequency specificity of lch5 neurons, we looked at the putative mechanotransduction channels that were reported to express specifically in subsets of lch5 neurons. Brv1 is a TRP channel that was reported to be relevant in temperature sensing and mechanosensation (25, 26). Previous study demonstrates that Brv1 is expressed in two neurons (neurons 1 and 3) in the lch5 (27) (Figure 3A). When Brv1 was knocked down in lch5, the response scores of 0.5 and 5 kHz sound stimulation dropped significantly (Figure 3B). Ca^2+^ responses of sub-neurons 1 and 3 to sounds were also reduced in Brv1-knocking down larvae (Figures 3C, S3B and S3C). Noticeably, the Ca^2+^ response of the 3^rd^ neuron did not completely disappear, suggesting the neuron’s sound sensing was only partially affected by Brv1. Thus, these results argue that Brv1 plays specific roles in detecting 0.5 and 5 kHz sounds.

**Figure 3.**
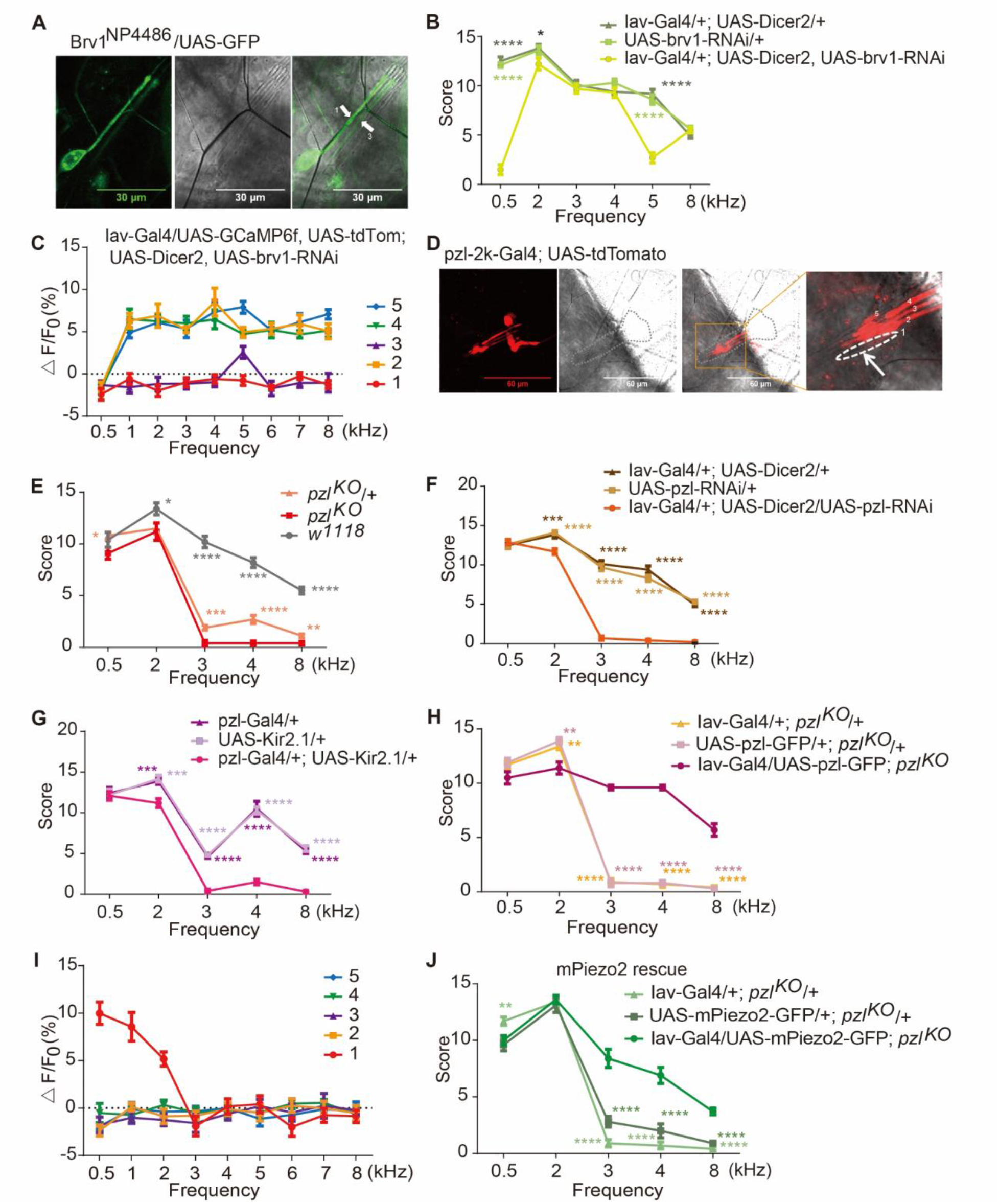
Brv1 and Pzl mediate hearing sensation to certain frequency ranges. See also Figure S3. (A) Brv1^NP4486^ labeled 2 lch5 neurons in *Drosophila* larvae. (B) Average sound response score for Brv1-knockdown larvae (Mean ± SEM, n=10; p*<0.05, p****<0.0001, one-way ANOWA with Dunnett’s multiple-comparisons test). (C) Statistical analysis of the Ca^2+^ responses of each lch5 neuron to different sound stimulations in Brv1-knockdown larvae (n=5). (D) pzl-Gal4 labeled 4 lch5 neurons in *Drosophila* larvae. (E) Average sound response score for *w*^*1118*^, *pzl*^*KO*^ and *pzl*^*KO*^/+ under different stimulation (n=10, Mean±SEM, p*<0.05, p**<0.01, p***<0.001, p****<0.0001, one-way ANOWA with Dunett’s multiple-comparisons test). (F) Average sound response score for Pzl-knockdown larvae (n=10, Mean±SEM, p***<0.001, p****<0.0001, one-way ANOWA with Dunett’s multiple-comparisons test). (G) Average sound response score when Pzl positive Cho neurons were blocked (n=10, Mean±SEM, p***<0.001, p****<0.0001, one-way ANOWA with Dunett’s multiple-comparisons test). (H) Average sound response score when Pzl was specifically rescued in Cho neurons of *pzl*^*KO*^ allele (n=10, Mean±SEM, p**<0.01, p****<0.0001, one-way ANOWA with Dunett’s multiple-comparisons test). (I) Statistical analysis of the Ca^2+^ responses of each lch5 neuron to different sound stimulations in *pzl*^*KO*^ allele (n=5). (J) Average sound response score when mouse Piezo2 was expressed in Cho neurons of *pzl*^*KO*^ allele (n=10, Mean±SEM, p**<0.01, p****<0.0001, one-way ANOWA with Dunett’s multiple-comparisons test).

Pzl, a gene that was found to be involved *Drosophila* larval locomotion (28), only expressed in neurons 2-5 of lch5 but not in neuron 1 (Figure 3D). This echoes the previous finding that Pzl is dispensable for low frequency (100 Hz-1,000 Hz) hearing (28). To explore whether Pzl plays a role in high frequency hearing, we first examined the behavioral response of *pzl*^*KO*^ larvae to 0.5, 2, 3, 4, and 8 kHz pure tones. We found that the response scores of *pzl*^*KO*^ larvae were significantly lower than those of wild type larvae to high frequency stimulations (greater than 3 kHz) (Figure 3E). Besides, larvae with either Pzl knocked-down (Iav > pzl-RNAi) specifically in Cho neurons or Pzl^+^ neurons blocked by Kir2.1 showed similar abnormal phenotypes in high frequency hearing (Figures 3F and 3G). Importantly, the behavioral defects in *pzl*^*KO*^ larvae were fully rescued by specifically expressing Pzl in Cho neurons (Figure 3H), further validating the role of *pzl* in mediating high frequency sound. We also found that only the 1^st^ lch5 neuron responded to lower frequency stimulations (0.5, 1 and 2 kHz) in *pzl*^*KO*^ larvae while the other neurons lost their sound sensitivity (Figure 3I). These results demonstrate that Pzl is required in high frequency sound sensation.

Previous study demonstrates that there is functional conservation between Pzl and Piezo family (28) which is essential for mechanotransduction across different species(29–35). Among them, mouse Piezo2 was found to be expressed in the hair cells (36). We found the hearing loss to high frequency in *pzl*^*KO*^ allele can be rescued by ectopically expressing mouse Piezo2 (mPiezo2) in Cho neurons (Figure 3J). We also tested fly DmPiezo and mammalian Piezo1 (human Piezo1 and mouse Piezo1). However, we found that none of the three proteins could fully rescue the behavioral defects in *pzl* mutant when expressed in Cho neurons (Figures S3E and S3F). Hence, mammalian Piezo2 protein and Pzl seem to share high functional homology in sensing high-frequency sound.

## Discussion

Here we have demonstrated that *Drosophila* larvae can sense a wide range frequencies of sound and the lch5 neurons mediate frequency discrimination. The rather simple architecture, and its genetic tractability, yet with functions in multiple sensory modalities make *Drosophila* lch5 organs prime paradigms for addressing central aspects of hearing transduction and tonotopical coding. Besides, our study provides an inroad to understand the ancestral origin of frequency heterogeneity of auditory sensory neurons.

### Tonotopy in the hearing organ of adult fly

In adult flies, Johnston’s organ in the secondary segment of the antenna is a specialized Cho with about 480 primary sensory neurons categorized into five subgroups with different functions (A-E)(6, 7, 37). Subgroups of Johnston’s organ are found to be frequency-dependent, with subgroup A mainly contributing to higher-frequency responses (100-1,000 Hz) and subgroup B displaying a clear preference for low-frequency vibrations (10-100 Hz)(7). Although they share similar developmental programs and structures with larval Cho, it’s not clear whether adult JO neurons can sense sounds in the kilo Hertz range. Nevertheless, Pzl protein only expresses in a subtype of JO neurons (Figure S3G), implicating those neurons may be preferentially tuned to high-frequency sound.

### Transduction mechanisms for sound in Cho

Three transient receptor potential (TRP) family cation channels have been implicated in mechano-electrical transduction in *Drosophila* chordotonal mechanoreceptors, i.e. NompC (TRPN1)(19) and the two vanillioid (TRPV) channels Nanchung (Nan) and Inactive (Iav)(12, 18). In Cho neurons, NompC localizes to the tips of the dendritic cilia, whereas Nan and Iav co-localize in the proximal cilium region(38, 39). NompC is a bona fide mechanotransduction channel(40), and Nan and Iav form a heteromeric channel whose activation mechanism is unknown. It’s still unclear how these channels orchestrate to transduce mechanical force. Nevertheless, the expression of the three channels appear homogenous in the five sub-neurons of lch5, so they are unlikely to contribute directly to the tonotopical gradient of the neurons. In contrast, Pzl and Brv1 seem to mediate the frequency selectivity, but whether they are part of the mechanotransduction machinery demands further investigation. Furthermore, Cho neurons are involved in a variety of biological processes in *Drosophila*. Besides hearing, they have been implicated in lower temperature sensation(41, 42), proprioception(22, 28, 42) and gentle touch response(42, 43). As a sensor with multifaceted roles, it’s an important question how the functional specificity is established.

### Central representation of Cho tonotopy

The neuronal circuits lch5 receptors feed into have recently been reconstructed at the electron microscopic level (44), setting the stage to analyze how sensory inputs from lch5 receptors are centrally processed. lch5 receptors reportedly convey mechanosensory information to central interneurons in the VNC (ventral nerve cord) that integrate mechanosensory and nociceptive inputs and whose activation causes stereotyped avoidance behaviors (45). Our results suggest that the downstream circuits are able to decode the combinatorial coding of frequency-specific activities among the five lch5 sub-neurons. How’s this achieved? One mechanism could be that low and high frequency neurons have different targets in the VNC, a labelled-line mechanism that is widely used by the somatosensory system (46).

### A conserved role in high-frequency hearing between Pzl and mammalian Piezo2

The mechanically gated ion channel family Piezo families play essential roles in diverse physiological processes (30, 47–53). There are two piezo genes in both human and mice. Here, our work demonstrates that mammalian Piezo2 could fully rescue high frequency hearing defects in *pzl* mutants but not mammalian Piezo1 or DmPiezo. It’s an interesting question what features of the channels confer their differential responsiveness (Pzl/Piezo2 vs. DmPiezo/Piezo1) to high frequency sound. Although the Piezo proteins exhibit high sequence conservation from fly to mammals in the transmembrane domains(28), structural studies demonstrate that Piezo1 and Piezo2 each has some unique motifs in spite of an overall similar architecture and the inactivation kinetics between Piezo1(slower) and Piezo2 (faster) are also distinct at negative membrane potentials (32, 54–57). Our results show that ectopically expressed Piezo2 rescues *pzl* mutant phenotype in hearing, arguing that the difference on frequency sensing may be molecularly intrinsic.

## Materials and methods

### Fly stocks and genetics

Pzl-RNAi flies were from Vienna *Drosophila* Stock Center (109995). Brv1-RNAi (31507), Iav-Gal4 and UAS-Kir2.1 were from Bloomington *Drosophila* Stock Center. Brv1^NP4486^ flies were from *Drosophila* Genetic Resource Consortium. UAS-hPiezo1 and UAS-mPiezo1 were gifts from Yuh-Nung Jan lab at UCSF. Flies were raised on standard medium at 25°C, 70 % humidity and 12 h/12 h light-dark cycles.

### Molecular cloning and generation of transgenic flies

UAS-mPiezo2-GFP was generated by inserting the coding sequence of mouse piezo2 into the pJFRC81 vector with GFP(58). Insertions were confirmed with PCR that detects UAS-mPiezo2-GFP sequence. Transgenic flies were generated by inserting the cassette to attp2 or 40 sites.

### Behavioral assay

Sound avoidance behavior was performed as previously described(59). Briefly, about 20 third instar larvae were rinsed with distilled water and transferred to the center of 1% agarose arena (a Petri dish with a diameter of 1500 mm). The arena was illuminated by 850 nm infrared LED lights and a speaker was placed at randomly locations along a concentric circle outside the arena. Larvae were put onto the central of the agarose arena before being videotaped with a camera (Basler GenlCam) placed atop the setup. The larvae were stimulated with different sound stimulation (a bird song, composite tone or pure tones) with the intensity ranged from 93 to 95 dB, SPL. After 90s stimulation, the distribution of the larvae was analyzed with ImageJ and the numbers of larvae on each side (referred to the speaker) were counted. The experiments were repeated at least for 3 times. Performance index was calculated as = (number of larvae on the near sound filed - number of larvae on the field far from the sound)/ total number of the larvae. The composited tone and pure tones were generated by MATLAB software and the pure tones were ramped to minimize acoustic onset/offset transients.

The responses of single third instar larva to sound were assessed modified according to previous study(12). The plate was placed beneath a speaker and the distance between the speaker and the plate was about 3 cm. The larvae were allowed to move freely for 1 min and were stimulated with a 1-s sound for 5 times with different frequencies (0.5, 2, 3, 4, 5 and 8 kHz) and an interval of 10 s. The sound was generated by a signal generator (RIGOL, DG1022U). The first behavior was counted when larva was stimulated according to the score we set: head lifting or turning for 1 point, pausing for 2 points and head contraction for 3 points.

### *In vivo* calcium imaging

*In vivo* calcium imaging of larval Cho neurons was performed with third instar larvae modified from previously study(12). Briefly, a freely moving larva was pressed between two coverslips with a drop of ddH2O to reduce its movement and the left lateral side of the larva is on top. The speaker was fixed by the side of microscope with its central point source straight to the larva. The sound intensity of the larva position was 94±1 dB, SPL. The imaging data were acquired with Olympus FV1000 confocal microscope when given sound stimulation (94±1 dB). The calcium indicator GCaMP6f was used to measure the Ca^2+^ signal, and the fluorescence intensity changes of tdTomato were used as controls. The GCaMP fluorescence demonstrated a dramatic increase on sound stimulation. A region of interest on each lch5 neuron was selected to measure GCaMP fluorescence intensity. The average GCaMP signal from the first 4 s before application of stimulus was taken as fluorescence F_0_, and ΔF/F_0_ was calculated for each data point. To ensure there is no artificial vibration, we conducted the non-contact vibration analysis to detect vibration frequency of the glass slide with a versatile laser vibrometer system (Polytec OFV-5000) under different frequencies of sound stimulation (1, 2, 3, 4, 5, 6, 7, 8 kHz, 94±1 dB, SPLs). The vibration frequency of the glass slide was same with the frequency of sound stimulation (Figures S3A and S3B), indicating that there is no other artificial vibration.

### Data analyses and statistical methods

The bird song was analyzed with MATLAB software. All data in bar and line graphs are expressed as Means ± SEMs. Student’s t test was used to evaluate the statistical significance between two datasets with Graphpad Prism 6 (Graphpad software Inc.). Statistical significance is indicated by ns for no significance, * for p<0.05, ** for p<0.01, *** for p<0.001 and **** for p<0.0001.

## Acknowledgments

We thank Dr. Wei Xiong for helping with sound stimulation. This work was supported by grant 31871059 from the National Natural Science Foundation of China, grant Z181100001518001 from the Beijing Municipal Science & Technology Commission, and a ‘Brain+X’ Seeds grant from the IDG/McGovern Institute for Brain Research at Tsinghua to W.Z. W.Z. is an awardee of the Young Thousand Talent Program of China.

## Author contributions

Y.H., Y.J., T.D., T.L., and W.Z. performed the experiments. Y.H. and W.Z. analyzed data and wrote the manuscript. All authors discussed and commented on the manuscript.

## Data and materials availability

All data is available in the main text and the supplementary materials.

## References

1. Z. F. Mann, M. W. Kelley, Development of tonotopy in the auditory periphery. Hearing Research 276, 2–15 (2011).

2. A. C. Mason, G. S. Pollack, “Introduction to Insect Acoustics” in Insect Hearing: With 53 Illustrations, G. S. Pollack, A. C. Mason, A. N. Popper, R. R. Fay, Eds. (2016), vol. 55, pp. 1–15.

3. G. S. Pollack, “Hearing for Defense” in Insect Hearing: With 53 Illustrations, G. S. Pollack, A. C. Mason, A. N. Popper, R. R. Fay, Eds. (2016), vol. 55, pp. 81–98.

4. A. Kamikouchi, Y. Ishikawa, “Hearing in Drosophila” in Insect Hearing: With 53 Illustrations, G. S. Pollack, A. C. Mason, A. N. Popper, R. R. Fay, Eds. (2016), vol. 55, pp. 239–262.

5. H. Hertweck, Anatomy and variability of the nervous system and the sense organs of Drosphila melanogaster (Meigen). Zeitschrift Fur Wissenschaftliche Zoologie 139, 559–663 (1931).

6. E. Matsuo et al., Identification of novel vibration- and deflection-sensitive neuronal subgroups in Johnston’s organ of the fruit fly. Frontiers in Physiology 5 (2014).

7. A. Kamikouchi et al., The neural basis of Drosophila gravity-sensing and hearing. Nature 458, 165–U161 (2009).

8. J. T. Albert, M. C. Goepfert, Hearing in Drosophila. Current Opinion in Neurobiology 34, 79–85 (2015).

9. Q. Hu et al., Edible bird’s nest enhances antioxidant capacity and increases lifespan in Drosophila Melanogaster. Cellular and Molecular Biology 62, 116–122 (2016).

10. R. S. Santoso, *Drosophila* melanogaster as a Feed Supplement for Swallow (*Collacalia fuchiphaga*). International Journal of Applied Chemistry 13, 89–97 (2017).

11. A. Novick, J. R. Vaisnys, ECHOLOCATION OF FLYING INSECTS BY BAT CHILONYCTERIS PARNELLII. Biological Bulletin 127, 478–& (1964).

12. W. Zhang, Z. Yan, L. Y. Jan, Y. N. Jan, Sound response mediated by the TRP channels NOMPC, NANCHUNG, and INACTIVE in chordotonal organs of Drosophila larvae. P Natl Acad Sci USA 110, 13612–13617 (2013).

13. D. J. Mennill et al., Wild Birds Learn Songs from Experimental Vocal Tutors. Current Biology 28, 3273–+ (2018).

14. I. Macova et al., Neurod1 Is Essential for the Primary Tonotopic Organization and Related Auditory Information Processing in the Midbrain. J Neurosci 39, 984–1004 (2019).

15. K. Karmakar et al., Hox2 Genes Are Required for Tonotopic Map Precision and Sound Discrimination in the Mouse Auditory Brainstem. Cell Reports 18, 185–197 (2017).

16. J. Kim et al., A TRPV family ion channel required for hearing in Drosophila. Nature 424, 81–84 (2003).

17. T. Voets, K. Talavera, G. Owsianik, B. Nilius, Sensing with TRP channels. Nature Chemical Biology 1, 85–92 (2005).

18. R. G. Walker, A. T. Willingham, C. S. Zuker, A Drosophila mechanosensory transduction channel. Science 287, 2229–2234 (2000).

19. S. Sidi, R. W. Friedrich, T. Nicolson, NompC TRP channel required for vertebrate sensory hair cell mechanotransduction. Science 301, 96–99 (2003).

20. Z. F. Gong et al., Two interdependent TRPV channel subunits, inactive and Nanchung, mediate hearing in Drosophila. J Neurosci 24, 9059–9066 (2004).

21. L. E. Cheng, W. Song, L. L. Looger, L. Y. Jan, Y. N. Jan, The Role of the TRP Channel NompC in Drosophila Larval and Adult Locomotion. Neuron 67, 373–380 (2010).

22. D. Zanini et al., Proprioceptive Opsin Functions in Drosophila Larval Locomotion. Neuron 98, 67–+ (2018).

23. T.-W. Chen et al., Ultrasensitive fluorescent proteins for imaging neuronal activity. Nature 499, 295–+ (2013).

24. P. M. Salvaterra, T. Kitamoto, Drosophila cholinergic neurons and processes visualized with Gal4/UAS-GFP. Gene Expression Patterns 1, 73–82 (2001).

25. M. Gallio, T. A. Ofstad, L. J. Macpherson, J. W. Wang, C. S. Zuker, The Coding of Temperature in the Drosophila Brain. Cell 144, 614–624 (2011).

26. M. Zhang et al., Brv1 Is Required for Drosophila Larvae to Sense Gentle Touch. Cell Reports 23, 23–31 (2018).

27. D. A. G. Sánchez (2018) Linking senses: the genetics of Drosophila larval chordotonal organs. (Georg-August-Universität Göttingen, Göttingen), p 152.

28. Y. Hu, Z. Wang, T. Liu, W. Zhang, Piezo-like Gene Regulates Locomotion in Drosophila Larvae. Cell Reports 26, 1369–+ (2019).

29. K. Nonomura et al., Piezo2 senses airway stretch and mediates lung inflation-induced apnoea. Nature 541, 176–+ (2017).

30. A. Faucherre, K. Kissa, J. Nargeot, M. E. Mangoni, C. Jopling, Piezo1 plays a role in erythrocyte volume homeostasis. Haematologica 99, 70–75 (2014).

31. S. E. Kim, B. Coste, A. Chadha, B. Cook, A. Patapoutian, The role of Drosophila Piezo in mechanical nociception. Nature 483, 209–212 (2012).

32. B. Coste et al., Piezo1 and Piezo2 Are Essential Components of Distinct Mechanically Activated Cation Channels. Science 330, 55–60 (2010).

33. S. Maksimovic et al., Epidermal Merkel cells are mechanosensory cells that tune mammalian touch receptors. Nature 509, 617–621 (2014).

34. E. R. Schneider et al., Neuronal mechanism for acute mechanosensitivity in tactile-foraging waterfowl. P Natl Acad Sci USA 111, 14941–14946 (2014).

35. K. Schrenk-Siemens et al., PIEZO2 is required for mechanotransduction in human stem cell-derived touch receptors. Nat Neurosci 18, 10–+ (2015).

36. Z. Wu et al., Mechanosensory hair cells express two molecularly distinct mechanotransduction channels. Nat Neurosci 20, 24–33 (2017).

37. Y. Ishikawa, N. Okamoto, M. Nakamura, H. Kim, A. Kamikouchi, Anatomic and Physiologic Heterogeneity of Subgroup-A Auditory Sensory Neurons in Fruit Flies. Frontiers in Neural Circuits 11 (2017).

38. J. Park et al., dTULP, the Drosophila melanogaster Homolog of Tubby, Regulates Transient Receptor Potential Channel Localization in Cilia. Plos Genetics 9 (2013).

39. C. Guan, N. Scholz, M. Nieberler, R. J. Kittel, T. Langenhan, Functional and genetic dissection of mechanosensory organs of Drosophila. Acta Physiologica 216 (2016).

40. P. Jin et al., Electron cryo-microscopy structure of the mechanotransduction channel NOMPC. Nature 547, 118–+ (2017).

41. Y. Kwon, W. L. Shen, H. S. Shim, C. Montell, Fine Thermotactic Discrimination between the Optimal and Slightly Cooler Temperatures via a TRPV Channel in Chordotonal Neurons. J Neurosci 30, 10465–10471 (2010).

42. J. C. Caldwell, M. M. Miller, S. Wing, D. R. Soll, D. F. Eberl, Dynamic analysis of larval locomotion in Drosophila chordotonal organ mutants. P Natl Acad Sci USA 100, 16053–16058 (2003).

43. N. Scholz et al., The Adhesion GPCR Latrophilin/CIRL Shapes Mechanosensation. Cell Reports 11, 866–874 (2015).

44. N. Scholz et al., Mechano-dependent signaling by Latrophilin/CIRL quenches cAMP in proprioceptive neurons. Elife 6 (2017).

45. T. Ohyama et al., High-Throughput Analysis of Stimulus-Evoked Behaviors in Drosophila Larva Reveals Multiple Modality-Specific Escape Strategies. Plos One 8 (2013).

46. J. C. Pereira, Jr., R. C. Alves, The labelled-lines principle of the somatosensory physiology might explain the phantom limb phenomenon. Medical Hypotheses 77, 853–856 (2011).

47. B. Coste et al., Gain-of-function mutations in the mechanically activated ion channel PIEZO2 cause a subtype of Distal Arthrogryposis. Proc Natl Acad Sci U S A 110, 4667–4672 (2013).

48. M. J. McMillin et al., Mutations in PIEZO2 Cause Gordon Syndrome, Marden-Walker Syndrome, and Distal Arthrogryposis Type 5. Am J Hum Genet 94, 734–744 (2014).

49. M. Okubo et al., A Family of Distal Arthrogryposis Type 5 Due to a Novel PIEZO2 Mutation. Am J Med Genet A 167, 1100–1106 (2015).

50. S. S. Ranade et al., Piezo2 is the major transducer of mechanical forces for touch sensation in mice. Nature 516, 121–U330 (2014).

51. S. M. Cahalan et al., Piezo1 links mechanical forces to red blood cell volume. Elife 4 (2015).

52. M. Pathak et al., The stretch-activated ion channel Piezo1 transduces matrix mechanics to direct human neural stem cell fate. Mol Biol Cell 25 (2014).

53. J. Wu, A. H. Lewis, J. Grandl, Touch, Tension, and Transduction - The Function and Regulation of Piezo Ion Channels. Trends Biochem Sci 42, 57–71 (2017).

54. J. Ge et al., Architecture of the mammalian mechanosensitive Piezol channel. Nature 527, 64–69 (2015).

55. Q. Zhao, H. Zhou, X. Li, B. Xiao, The mechanosensitive Piezo1 channel: a three-bladed propeller-like structure and a lever-like mechanogating mechanism. Febs Journal 286, 2461–2470 (2019).

56. Q. Zhao et al., Structure and mechanogating mechanism of the Piezol channel. Nature 554, 487–+ (2018).

57. L. Wang et al., Structure and mechanogating of the mammalian tactile channel PIEZO2. Nature 573, 225–+ (2019).

58. B. D. Pfeiffer, J. W. Truman, G. M. Rubin, Using translational enhancers to increase transgene expression in Drosophila. P Natl Acad Sci USA 109, 6626–6631 (2012).

59. S. Hueckesfeld, M. Peters, M. J. Pankratz, Central relay of bitter taste to the protocerebrum by peptidergic interneurons in the Drosophila brain. Nature Communications 7 (2016).

