## supplemental information for "Molecular Mechanisms for Frequency Specificity in a *Drosophila* Hearing Organ"

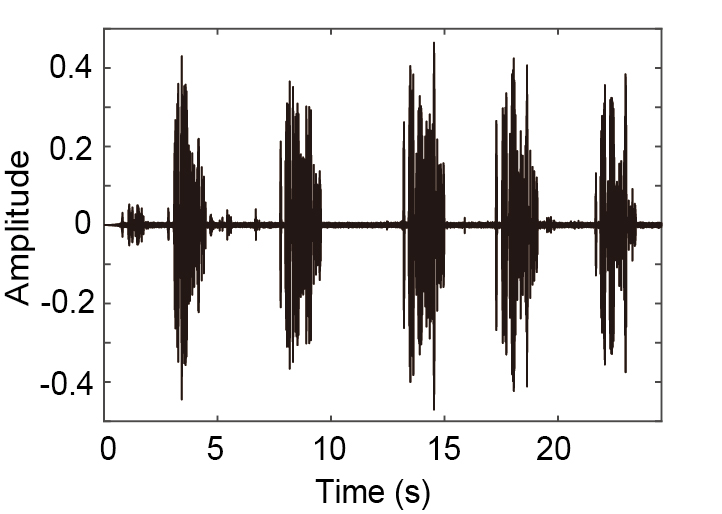

**Figure S1.** Spectrogram for the bird song

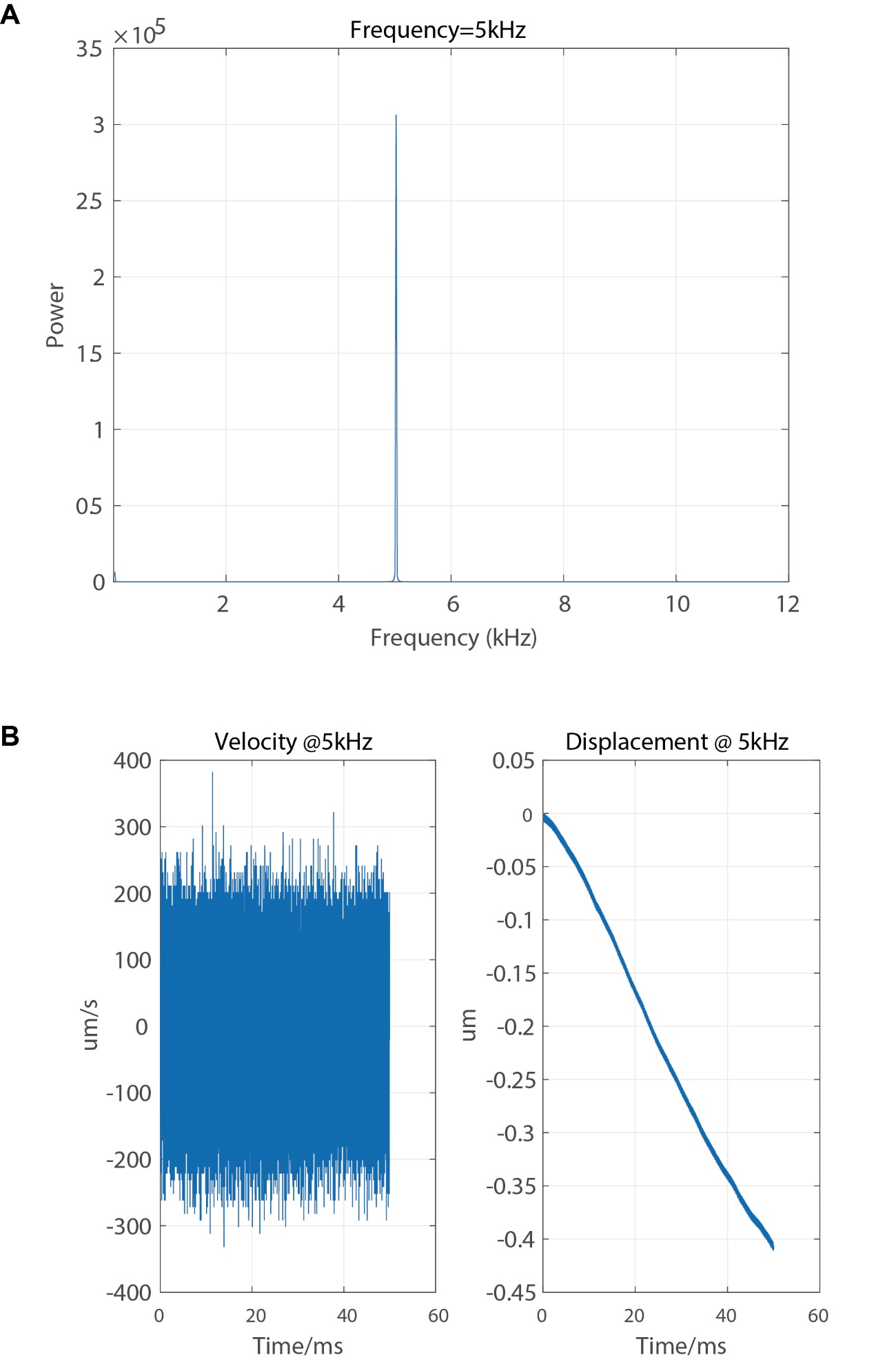

**Figure S2.** Non-contact vibration analysis

(A) An example of the glass slide frequency when stimulated with 5 kHz pure sound stimulation (94 ±1 dB, SPL).
(B) Left: velocity of the glass slide when stimulated with 5 kHz pure sound stimulation (94 ±1 dB, SPL); Right: displacement of the glass slide when stimulated with 5 kHz pure sound stimulation (94 ±1 dB, SPL).

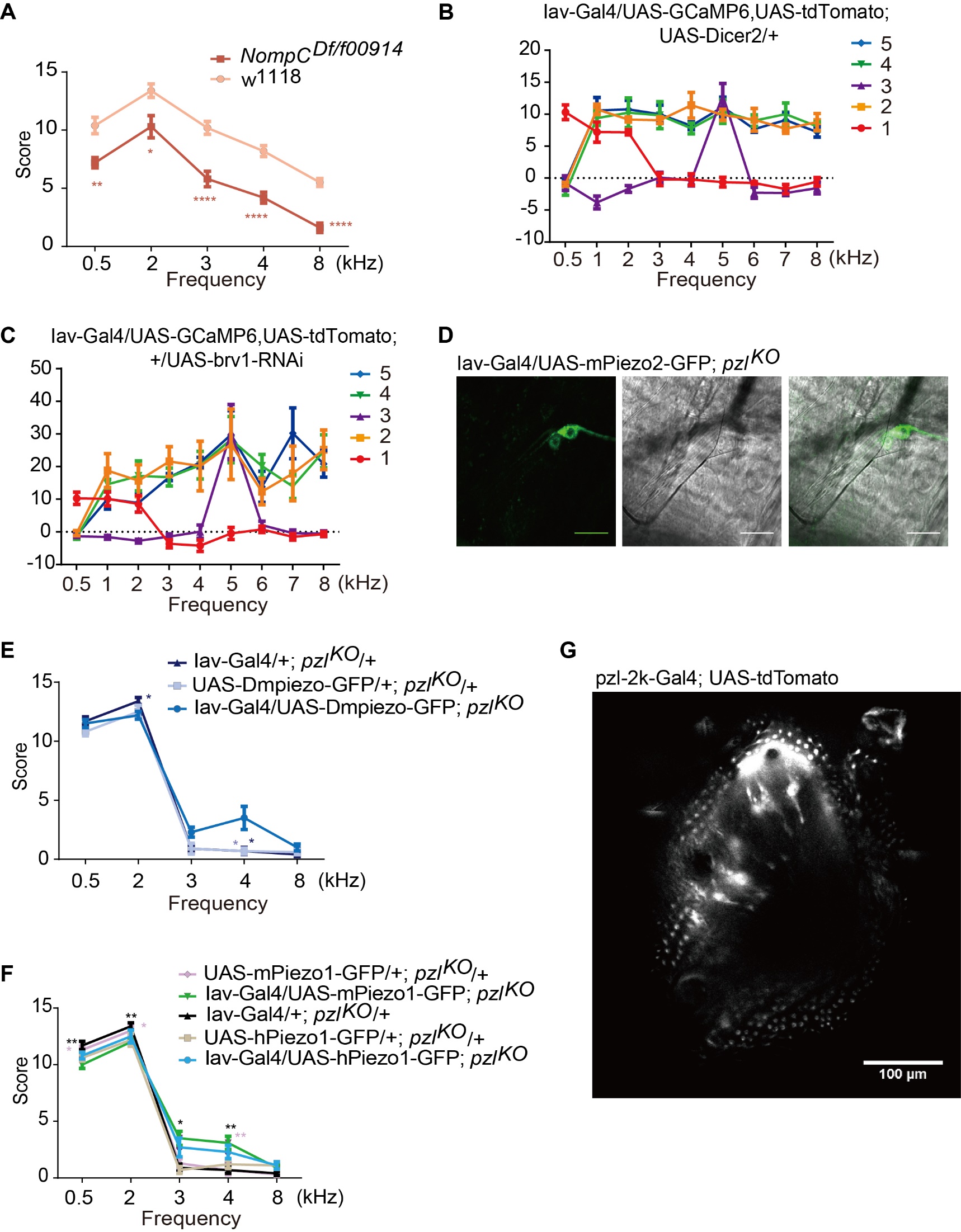

**Figure S3.** Brv1 and Pzl play distinct roles in hearing

(A) *NompC* mutant larvae show hearing defects both to low and high frequency sound stimuli. Mean ± SEM, n=10; * for p<0.05, ** for p<0.01 and **** for p<0.0001, unpaired t test.
(B) Statistical analysis of the Ca^2+^ responses of each lch5 neuron to different sound stimulations in control larvae (Iav-Gal4/UAS-GCaMP6, UAS-tdTomato; UAS-Dicer2/+, n=5).
(C) Statistical analysis of the Ca^2+^ responses of each lch5 neuron to different sound stimulations in control larvae (Iav-Gal4/UAS-GCaMP6, UAS-tdTomato; +/UAS-brv1-RNAi, n=5)
(D) Mouse Piezo2 is specifically expressed in larval chordotonal neurons.
(E) Average sound response score when *Drosophila* Piezo (DmPiezo) was expressed in Cho neurons of *pzl^KO^* allele (Mean ± SEM, n=10, p*<0.05, one-way ANOWA with Dunnett’s multiple-comparisons test).
(F) Average sound response score when human Piezo1 (hPiezo1) and mouse Piezo1 (mPiezo1) were expressed in Cho neurons of *pzl^KO^* allele (Mean ± SEM, n=10, *p<0.05, **p<0.01, one-way ANOWA with Dunnett’s multiple-comparisons test).
(G) Pzl is expressed in a subtype of JO neurons.

**Movie S1.** The third instar larva showed head contraction, head lifting and turning when given the bird song stimulation (94 dB, SPL).

**Audio S1.** A bird song of a meadow bunting (Emberiza cioides) which is widely spread in eastern Asia.
